# Striatal ensemble activity in an innate naturalistic behavior

**DOI:** 10.1101/2023.02.23.529669

**Authors:** Samuel Minkowicz, Mychaela Alexandria Mathews, Felicia Hoilam Mou, Hyoseo Yoon, Sara Nicole Freda, Ethan S Cui, Ann Kennedy, Yevgenia Kozorovitskiy

## Abstract

Self-grooming is an innate, naturalistic behavior found in a wide variety of organisms. The control of rodent grooming has been shown to be mediated by the dorsolateral striatum through lesion studies and in-vivo extracellular recordings. Yet, it is unclear how populations of neurons in the striatum encode grooming. We recorded single-unit extracellular activity from populations of neurons in freely moving mice and developed a semi-automated approach to detect self-grooming events from 117 hours of simultaneous multi-camera video recordings of mouse behavior. We first characterized the grooming transition-aligned response profiles of striatal projection neuron and fast spiking interneuron single units. We identified striatal ensembles whose units were more strongly correlated during grooming than during the entire session. These ensembles display varied grooming responses, including transient changes around grooming transitions or sustained changes in activity throughout the duration of grooming. Neural trajectories computed from the identified ensembles retain the grooming related dynamics present in trajectories computed from all units in the session. These results elaborate striatal function in rodent self-grooming and demonstrate that striatal grooming-related activity is organized within functional ensembles, improving our understanding of how the striatum guides action selection in a naturalistic behavior.

## Introduction

Self-grooming is an evolutionarily conserved, ethologically relevant, and innate behavior. Self-grooming is found in arthropods^1,2^, birds^3–5^, and mammals^6–10^. In mammals, grooming serves to care for the outer body surface, for de-arousal^11,12^, thermoregulation^13–15^, and water economy^16^. Rodent grooming is an innate behavior^9,17–20^, with mice showing primitive forms of facial grooming as early as postnatal day 1 (p1)^17^. From p4-8, the kinematics of grooming movements become more tightly coordinated. After p9, mice start to perform movements similar to those performed by adults. Rat pups begin to exhibit functional face grooming at p6-8^9^, but do not perform long sequences of grooming until p14-21^18–20^, the time of development when many complex motor programs begin to emerge^21,22^.

The selection and initiation of actions is controlled by the striatum^23–29^. The striatum has been shown to encode action space^30^ and to flexibly combine behavioral motifs into actions^25^. Cells within the striatum, predominantly consisting of striatal spiny projection neurons^31–33^, are organized into functional clusters of co-active units; this clustering is considered to be important for striatal network dynamics^34,35^ and behavioral control^30,36–40^. The striatum has been implicated in the production of self-grooming behavior, and lesions of the striatum disrupt grooming bouts^41^. Within the striatum, neurons of the dorsolateral striatum^42,43^ and nearby central striatal regions^44^ display grooming-related activity. Yet, how populations or ensembles of neurons in the striatum—defined here as sets of neurons that are more likely than chance to be co-active—encode self-grooming remains unclear. Given the likely importance of ensemble activity in the striatum, elaborating whether grooming-associated neural activity maps onto striatal ensembles is significant, and it remains to be elucidated.

Here we recorded simultaneous activity of populations of neurons in the dorsolateral striatum of freely moving mice during spontaneous grooming using extracellular probes. Because grooming can be a relatively rare behavior, many hours of data must be acquired to capture an adequate sample of its neural correlates. To overcome this obstacle, we developed a semi-automated approach to detect grooming bouts from behavioral videos using 3D pose estimation and postural heuristics. We found striatal projection neurons (SPN) and fast spiking interneurons (FSI) with temporally diverse grooming-related activity. Furthermore, we identified striatal ensembles that encode core parameters of grooming bouts, including the transitions in and out of individual bouts, as well as bout duration.

## Results

Mice were placed in a transparent, triangular arena with video recorded from each side to capture spontaneous behavior (**Figure 1A, B**). We identified mouse grooming bouts in a semi-automated 4-step process (**Figure 1C**). First, mouse limb positions were tracked in 2D in each view using DeepLabCut and triangulated to 3D using Anipose (see Methods). The 3D limb positions were then used to isolate likely grooming times, using a set of postural heuristics including movement speed, whether the animal was rearing, and the hand-to-nose distance. These heuristics identified general windows during which grooming was likely to be occurring, but they did not capture the precise timing of the behavior. We therefore refined heuristic output using manual frame-by-frame annotation, to capture the precise start and stop times of grooming (**Figure 1—figure supplement 1A**). In 117 total hours of analyzed video, we observed 304.8 minutes of grooming behavior within 1,475 individual bouts of grooming; mice groomed for 4.1% of each session on average (**Figure 1D**, 4.1 ± 0.3%, 63 sessions from 6 mice). Grooming bouts were on average 12.4 seconds long (**Figure 1E**, 12.4 ± 0.4 seconds) and separated by 4.3 minutes (**Figure 1F**, 4.3 ± 0.1 minutes).

**Figure 1.**
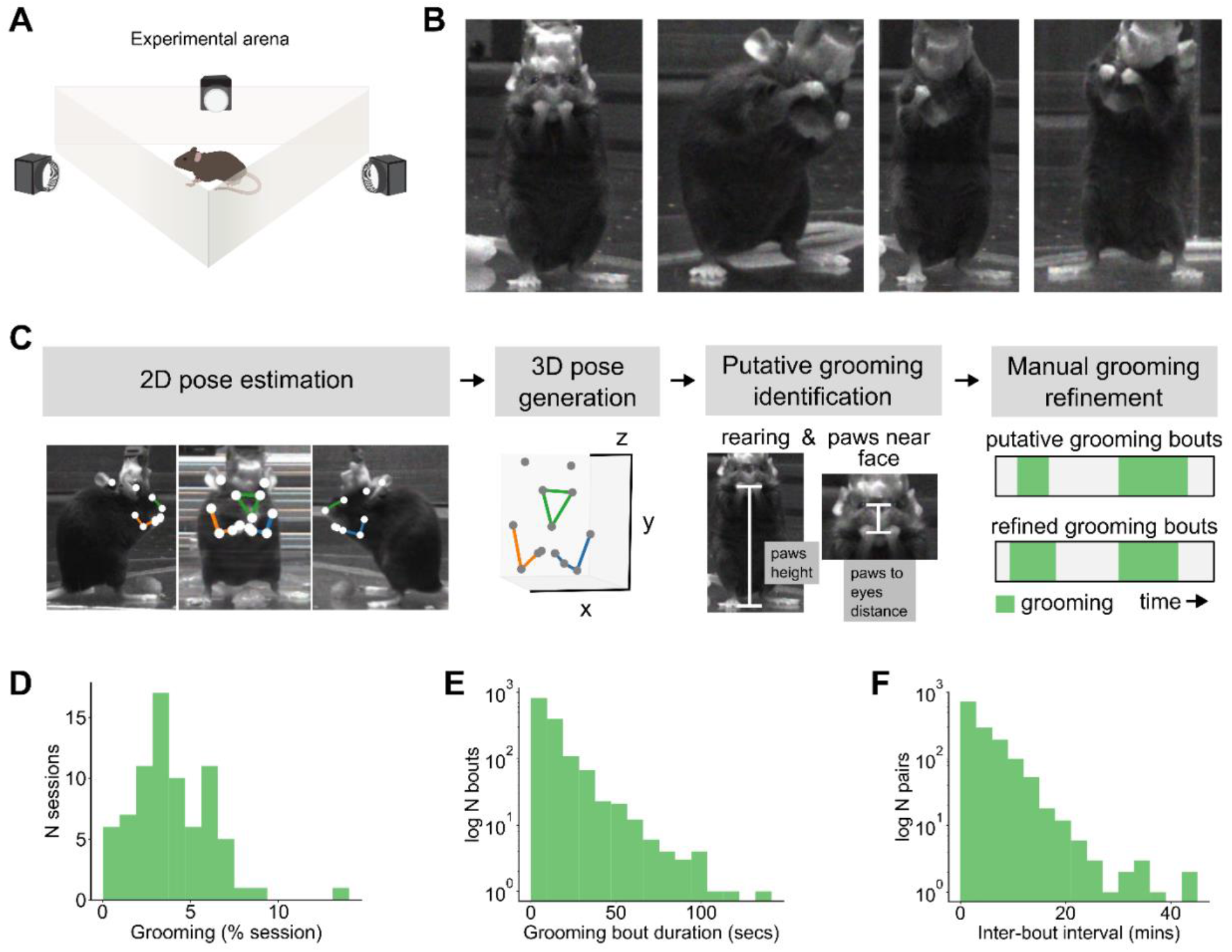
3D tracking and characterization of the structure of mouse spontaneous grooming. **A**. Schematic of experimental setup. Mice were placed into an equilateral triangular arena made of transparent acrylic (12-inch sides and height) and their behavior was captured with three side view cameras. Schematic is not to scale. **B**. Example video frames of mice during spontaneous grooming. **C**. Overview of our 4-step grooming identification approach. **D**. Distribution of the percent of each session that mice spent spontaneously grooming (6 mice, 63 sessions, 117 experiment hours, 1,475 grooming bouts, 304.8 total minutes of spontaneous grooming). **E**. Distribution of spontaneous grooming bout durations in seconds (for the same dataset as in **D**). **F**. Distribution of inter-bout intervals in minutes (for the same dataset as in **D**).

**Figure 1–figure supplement 1.**
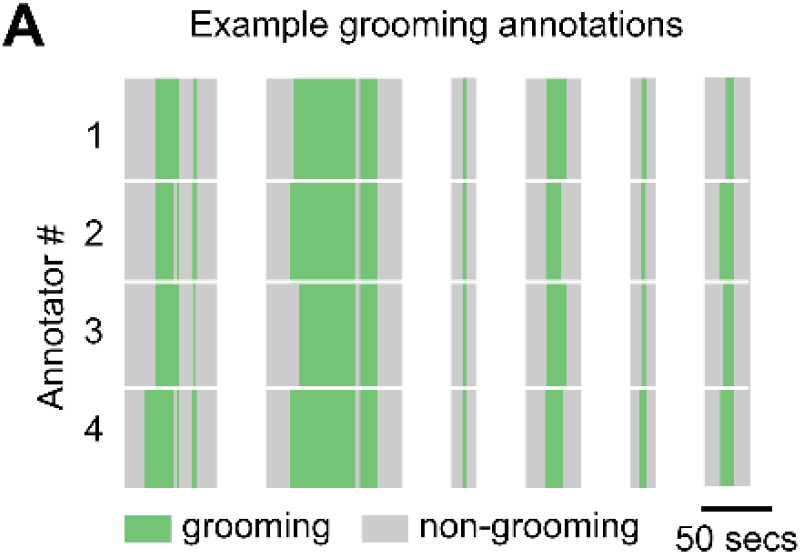
Manual refinement of grooming identification. **A**. Example annotations of grooming behavior by four annotators. Green regions in each row denote times that the annotator labeled as grooming and gray regions denote times not annotated as grooming. Vertically aligned white spaces indicate breaks of variable time between individual grooming bouts. Average pairwise Jaccard Index (or Intersection over Union) computed on all annotations from a single 2-hour session was 0.76 ± 0.04.

To record striatal activity during mouse spontaneous grooming, we implanted 64-channel electrodes into the dorsolateral striatum of adult male and female mice (**Figure 2—figure supplement 1A-D**, N=6 mice). We classified well-isolated units as putative striatal spiny projection neurons (SPN) and putative fast spiking interneurons (FSI) on the basis of their firing rates and spike waveform, following established criteria^45–47.^ Units that did not fit the criteria for SPNs or FSIs were labelled as ‘other’ and excluded from further analyses. (**Figure 2A**, SPN: 88.2% 2,755 units, FSI: 3.2% 100 units, other: 8.6% 269 units).

**Figure 2.**
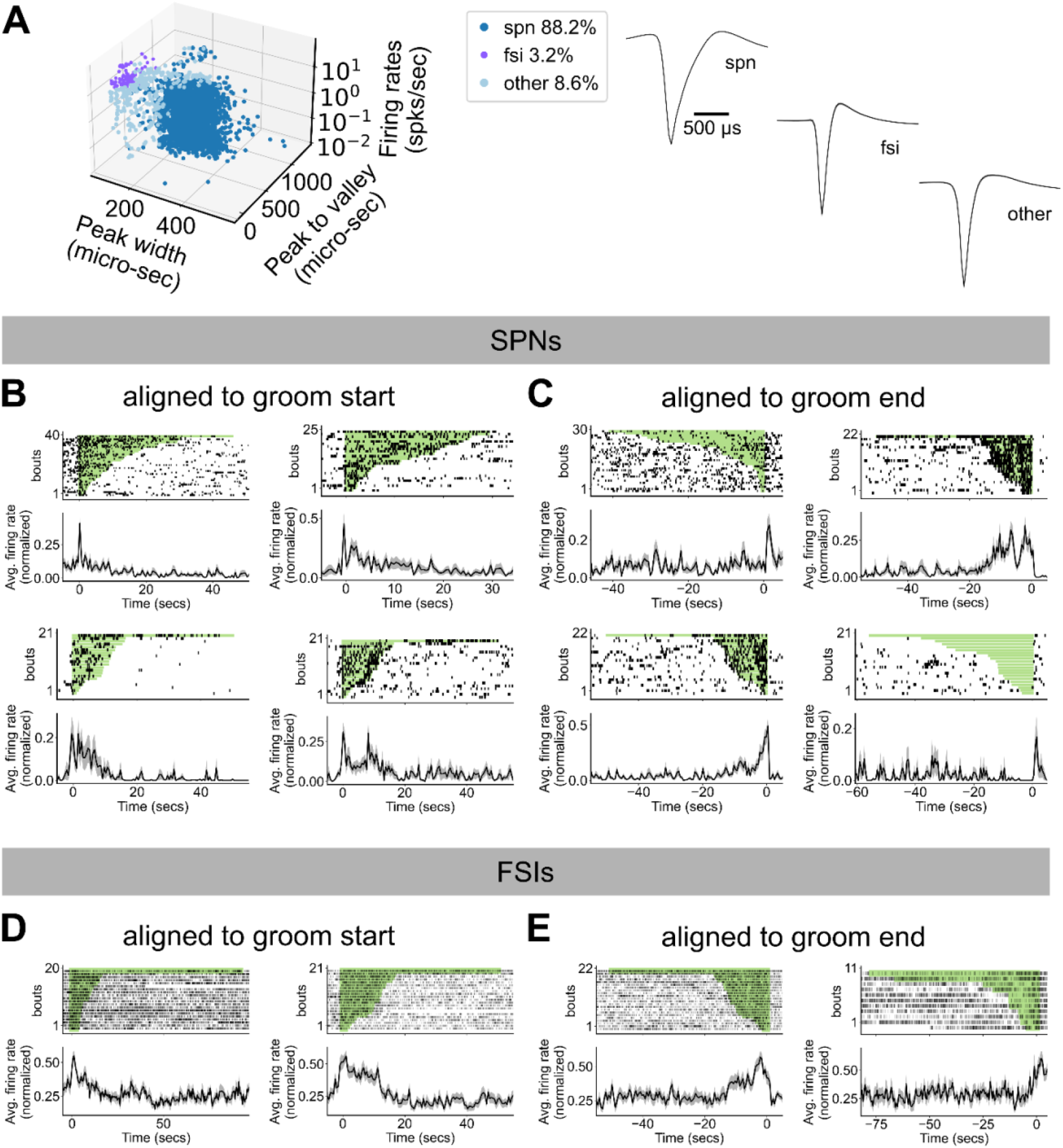
Striatal cell type specific activity mapped to spontaneous self-grooming. **A**. Classification of recorded units. Left: units were categorized as either SPNs, FSIs, or ‘other’ by their firing rates, spike waveform peak width, and duration between spike waveform peak to valley (SPN: 88.2% 2,755 units, FSI: 3.2% 100 units, other: 8.6% 269 units). Right: average +/- SEM spike waveforms for units in each category. **B**. Activity of 4 example SPNs during grooming. Neural activity are aligned to the start of grooming bouts, denoted by t = 0. Top: spike raster plots for the example neuron during each grooming bout in the given session. Grooming bouts are sorted by grooming bout duration, denoted by the green rectangles. Bottom: unit average firing rate aligned to grooming start and normalized to [0, 1]. **C**. Activity of 4 example SPNs as in **B**, but for aligned to groom end. **D**. Activity of 4 example units as in **B**, but for FSIs aligned to groom start. **E**. Activity of 4 example units as in **B**, but for FSIs aligned to groom end.

**Figure 2–figure supplement 1.**
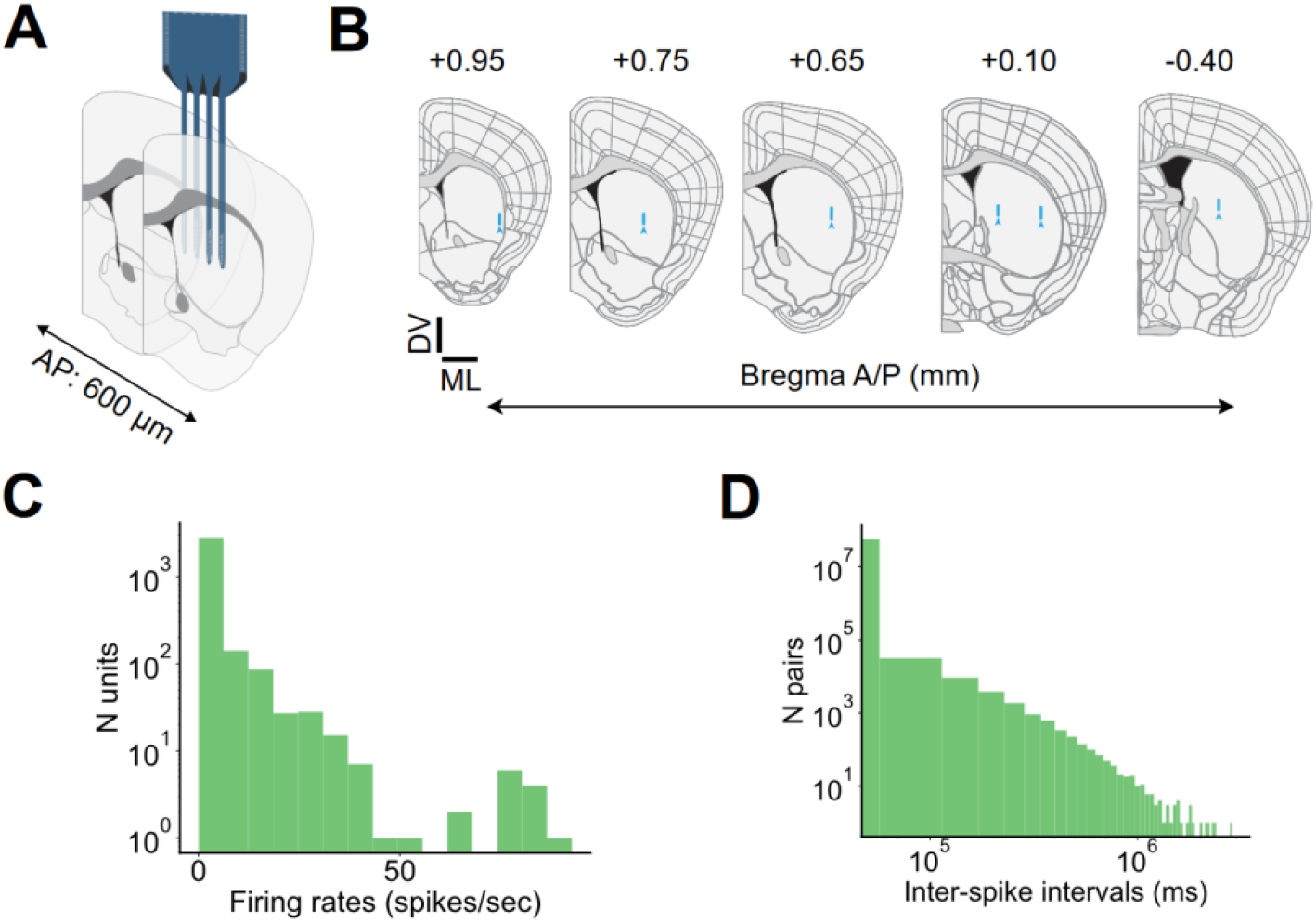
Probe placement and firing rate characteristics. **A**. Schematic depicting how the four shanks within our recording electrodes were implanted along the anterior-posterior axis (shanks span 600 μm). **B**. Representative coronal slices of electrode placements. Coronal histological slices were obtained and registered to the Allen Common Coordinate Framework (CCFv3) using the WholeBrain software package. Blue arrows depict the most ventral electrode position for each mouse. Blue lines depict the region covered by sites in one shank of each electrode. Anterior-Posterior (A/P) coordinates from Bregma used for registration are depicted above each slice. (6 mice, scale bar DV and ML = 1 mm). **C**. Distribution of firing rates for all recorded units (3,124 units. 6 mice). **D**. Distribution of inter-spike intervals for all recorded units (3,124 units. 6 mice).

To identify example individual units with grooming-related activity, we isolated units whose activity within a two-second window around the start or end of grooming was two standard deviations above mean activity during a grooming-free baseline period. All subsequent analyses were performed on the full dataset. Example SPNs and FSIs with grooming related activity are shown in **Figure 2B-E**. We observed units displaying diverse grooming-related responses, including increased activity at the start of grooming, end of grooming, and both start and end of grooming, as well as units that showed elevated activity for the duration of grooming, and units that showed reduced activity for the duration of grooming.

To characterize different grooming-associated responses in the recorded striatal units, we first performed principal components analysis (PCA)^48^ on the grooming transition-triggered-average activity of all recorded SPNs and FSIs. We performed PCA separately on SPN activity aligned to groom start, SPN activity aligned to groom end, FSI activity aligned to groom start, and FSI activity aligned to groom end. The first principal component (PC) from each group reflects units that undergo a step-like increase or decrease in their activity at the grooming transition (**Figure 3A, B, F, G, K, L, P, Q**, SPNs aligned to groom start: 336 units with positive contributions to the first PC (denoted “> 0”), 640 < 0, SPNs aligned to groom end: 526 > 0, 424 < 0, FSIs aligned to groom start: 66 > 0, 1 < 0, FSIs aligned to groom end: 22 > 0, 44 < 0). The second PC from each group reflects units whose activity transiently peaks or decreases around the grooming transition (**Figure 3C, D, H, I, M, N, R, S**, SPNs aligned to groom start: 364 > 0, 292 < 0, SPNs aligned to groom end: 164 > 0, 518 < 0, FSIs aligned to groom start: 4 > 0, 19 < 0, FSIs aligned to groom end: 5 > 0, 19 < 0). Taken together, these results show that single unit SPNs and FSIs encode transitions into and out of an innate naturalistic behavior.

**Figure 3.**
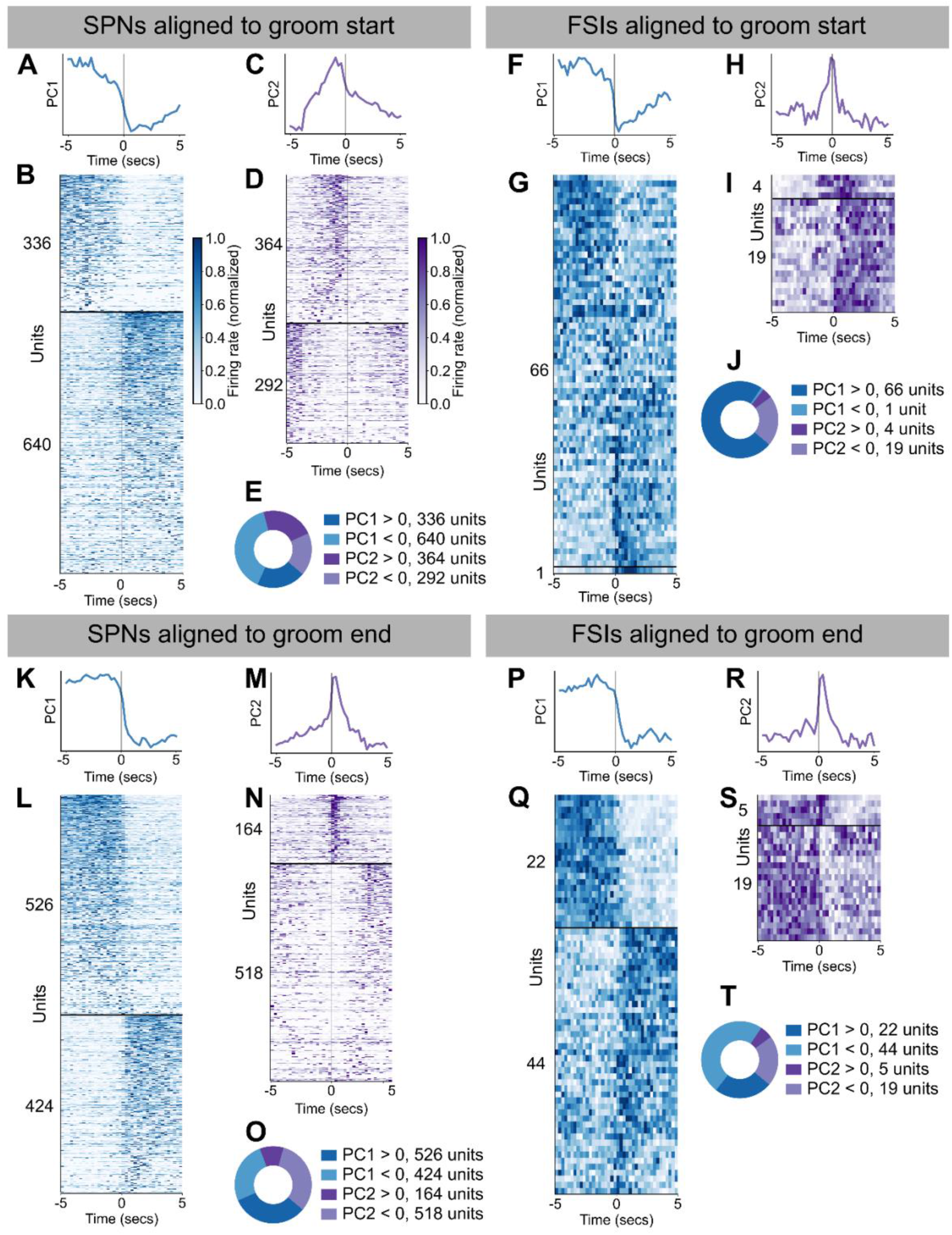
Emergent motifs in SPN and FSI activity around grooming transitions. **A**. First principal component from decomposing 10 seconds of SPN activity centered around grooming bout start times (1,632 units, explains 9.3% of variance). **B**. Activity around grooming start for units that had the largest magnitude weight for the first principal component. Units with weight > 0 are shown above the horizontal black line (336 units) followed by units with weight < 0 (640 units). Units are sorted by their weight for PC1. Each unit’s activity was normalized to the range from zero to one. **C**. Same as **A**, but for the second principal component (explains 5.1% of variance). **D**. Same as **B**, but for the second principal component (364 units with weight > 0, 292 units with weight < 0). **E**. Donut plot depicting the number of units with positive and negative weights for the first two principal components. **F-J**. Same as **A-E**, but for FSIs aligned to groom start (66 units with PC1 weight > 0, 1 unit with PC1 weight < 0, 2 units with PC2 weight > 0, and 19 units with PC2 weight < 0. PC1 explains 21.2% of variance and PC2 explains 7.1% of the variance). **K-O**. Same as **A-E**, but for SPNs aligned to groom end (526 units with PC1 weight > 0, 424 units with PC1 weight < 0, 164 units for PC2 weight > 0, and 518 units with PC2 weight < 0. PC1 explains 12.9% of variance and PC2 explains 5.2% of the variance). **P-T**. Same as **A-E**, but for FSIs aligned to groom end (22 units with PC1 weight > 0, 44 units with PC1 weight < 0, 5 units with PC2 weight > 0, and 19 units with PC2 weight < 0. PC1 explains 34.6% of variance and PC2 explains 8.6% of the variance).

The event-triggered average responses of striatal neurons reveal several patterns of firing rate modulation aligned with the start or end of grooming. However, because this analysis relies on trial-averaging, it cannot be used to determine whether the striatum contains ensembles of neurons that are consistently co-active. To address this question, we looked for functional clusters, or ‘ensembles’ of co-active striatal neurons with grooming related activity. To achieve this, we first constructed a matrix containing the grooming activity of all units in a given session, where the activity of each unit plus 5 seconds before and after grooming was concatenated. This matrix is shown for an example session in **Figure 4A**; dendrogram-based sorting of recorded units makes evident the presence of synchronous activity in groups of units. Synchrony among units is also suggested by the block-diagonal structure of the correlation matrix for this example session (**Figure 4B**). This synchrony is further suggested by comparison to a time-shuffled version of the data (**Figure 4—figure supplement 1A)** and its corresponding correlation matrix (**Figure 4—figure supplement 1B)**.

**Figure 4.**
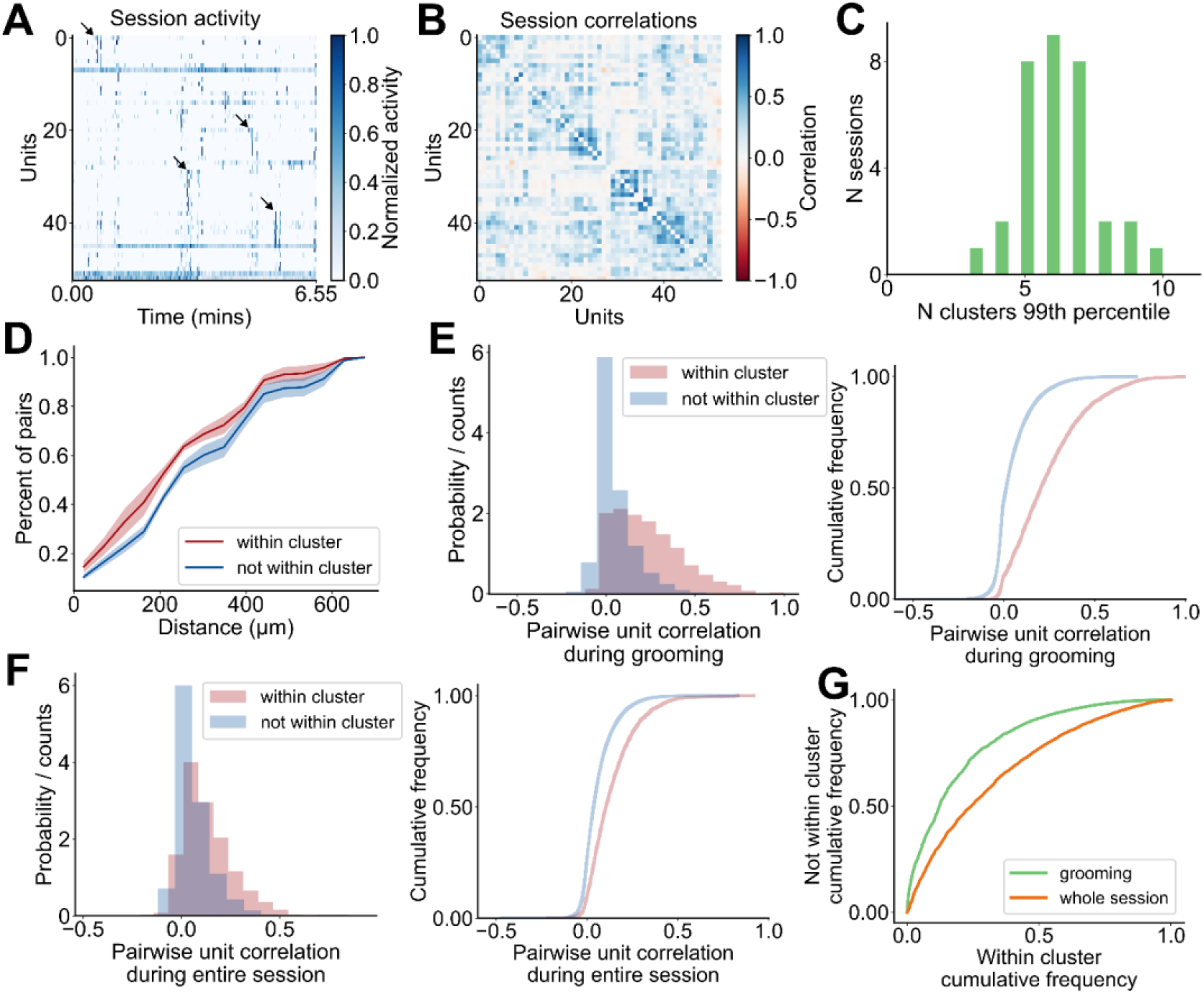
Identification and characterization of striatal grooming ensembles. **A**. Heatmap of activity during grooming for all units in an example session (53 units, 6.55 minutes). Arrows point out a subset of synchronous events. Each unit’s activity is normalized to [0, 1]. **B**. Correlation matrix for the activity shown in **A. C**. Distribution of statistical estimate for the number of ensembles within a given session (4 mice, 33 sessions). The estimate is obtained by computing the number of eigenvalues from the data that are above the 99^th^ percentile of the distribution of eigenvalues from 5,000 random shuffles of the data. **D**. Average cumulative distribution of pairwise unit distances for each pair of units that are within the same cluster (red) and each pair of units that are not within the same cluster (blue). Plot depicts the mean ± SEM of the cumulative distribution across mice (4 mice). **E**. Distribution of pairwise unit correlations during grooming for each pair of units that are within the same cluster (pink 2,747 pairs) and each pair of units that are not within the same cluster (blue 27,421 pairs). Left: histogram. Right: cumulative distribution. **F**. Same as in **E**, but for pairwise correlations computed from activity during the entire session including grooming times. **G**. Comparison between the difference in pairwise unit correlations for units within and not within the same cluster computed during grooming (green) and during the whole session (orange) (grooming AUC: 0.81, whole session AUC: 0.69).

To test for the presence of ensembles and to estimate the number of potential ensembles within each session, we used an eigenvalue-based statistical method^49,50^, where we identify dimensions of the neural covariance matrix that capture more variance than expected by chance and found each session to have three to ten ensembles (**Figure 4C**, 6.2 ± 0.3 ensembles, 33 sessions, see Methods for details). Next, using the meta-*k*-means clustering algorithm^51^, we identified the ensembles present in each session. Meta-*k*-means is an extension of *k*-means where one performs N iterations of *k*-means, keeping track of how many times each pair of units gets clustered together. The proportion of times that each pair gets clustered together is used to assign intermediate clusters before a final merging step. We chose to use meta-*k*-means for two reasons. First, meta-*k*-means does not always result in cluster assignment for all units, which is preferable because our recording configuration only samples a small subset of neurons in striatum and we do not expect all recorded units to belong to an identified striatal ensemble. Second, meta-*k*-means allows for the final number of identified ensembles to be greater or less than the initial choice of *k*, which is preferable because we do not have *a priori* knowledge of ensemble numbers present in each session.

We found that 80% of recorded SPNs (1,074 units) and 70% of recorded FSIs (41 units) were clustered into ensembles of two or more units **(Figure 4—figure supplement 1C**). The majority of clusters were composed of only SPNs, and FSIs were almost always assigned clusters together with SPNs (**Figure 4— figure supplement 1D;** 88% (214 clusters) SPN only, 1% (1 cluster) FSIs only, 11% (28 clusters) SPN and FSIs). Within our identified ensembles, units in the same ensemble tended to be spatially closer than units that are not within the same ensemble (**Figure 4D)**, consistent with striatal ensembles detected using imaging techniques^30,38,39^. During grooming, the pairwise correlation between units within the same cluster is higher than for the correlation between units not in the same cluster, whether those units are unclustered or belong to a different cluster (**Figure 4E**). The distributions of pairwise correlations computed using data from the whole session are shown in **Figure 4F**. This in-versus-out of cluster difference is weaker when comparing unit activity during the whole session, rather than during grooming (**Figure 4G**, grooming AUC: 0.81, whole session AUC: 0.69).

The distribution of the number of ensembles identified in each session is shown in **Figure 4—figure supplement 1E**. We found a significant positive relationship between the number of ensembles identified and the number of units in each session (**Figure 4—figure supplement 1F**, R^2^ = 0.239, p = 0.004). The distribution of the number of units within each cluster is shown in **Figure 4—figure supplement 1G** (median cluster size = 4 units, mean = 4.6 ± 0.2 units). Cluster size did not increase with the number of units recorded in a given session (**Figure 4—figure supplement 1H**, R^2^ = 0.01, p = 0.582). The distribution of the percent of clustered units in each session is shown in **Figure 4—figure supplement 1I** with half of sessions having at least 82% clustered units.

**Figure 4–figure supplement 1.**
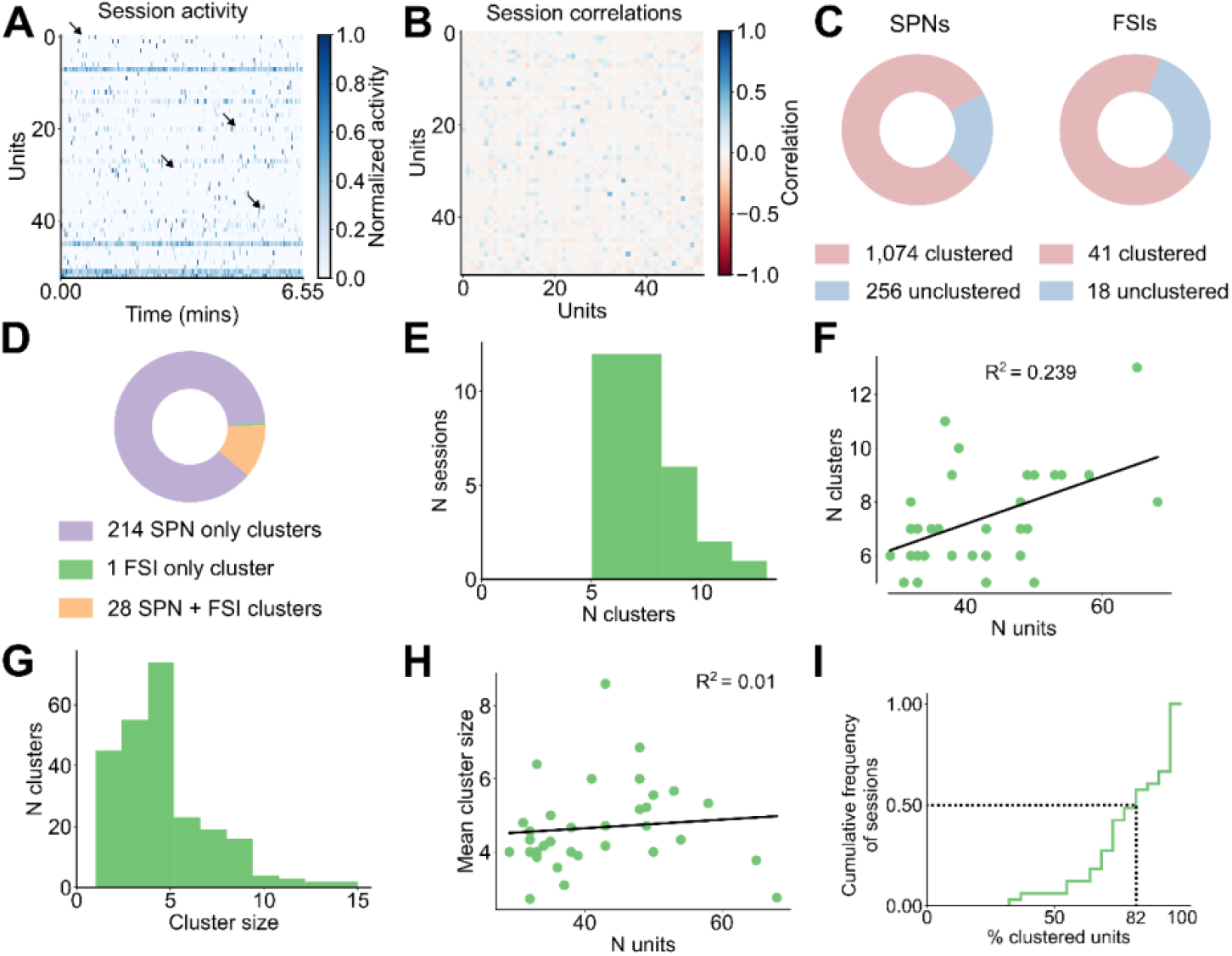
Statistics of identified clusters and activity patterns. **A**. Heatmap of activity during grooming with shuffled time bins for all units in the example session shown in **4A** (53 units, 6.55 minutes). Arrows are in the same position as in **Figure 4A** highlighting the absence of synchrony after shuffling. Each unit’s activity is normalized to the range from zero to one. **B**. Correlation matrix for the shuffled activity shown in **A. C**. Total number of SPNs and FSIs that were clustered (pink) and unclustered (blue) (4 mice, 33 sessions, 243 clusters). **D**. Total number of clusters that comprised of only SPNs (purple), only FSIs (green), and of both SPNs and FSIs (orange) (data as in **C**). **E**. Distribution of the number of clusters in each session. **F**. Relationship between the number of clusters found in each session and the number of units in that session (R^2^ = 0.239, p = 0.004). G. Distribution of the number of units in each cluster (data as in **C**). **H**. Relationship between the average cluster size in each session and the number of units in that session (R^2^ = 0.01, p = 0.582). **I**. Cumulative distribution of the percent of clustered units in each session. Dashed lines denote that half the sessions have at least 82% clustered units (data as in **C**).

Having demonstrated the existence of ensemble activity during grooming, we next asked whether ensemble activity was enriched during specific timepoints during grooming. A heatmap displaying the average grooming activity for all significant ensembles is shown in **Figure 5A** (112 ensembles, 4 mice, 33 sessions). To visualize the average grooming activity of ensembles across grooming bouts of varying duration, we first linearly time-warped ensemble activity during grooming. We found that ensemble peak activity is enriched around the transitions into and out of grooming (**Figure 5B**), but that the dynamics of grooming ensemble activation are diverse. For example, we found ensembles with higher activity during grooming, lower activity during grooming, and with peak activity at the start or end of grooming, with individual cluster examples of these patterns shown in **Figure 5 C-F** and **Figure 5—figure supplement 1C**. This finding was unchanged when we analyzed the percentage of active units within each ensemble rather than the ensemble-average firing rate (**Figure 5—figure supplement 1A, B**).

**Figure 5.**
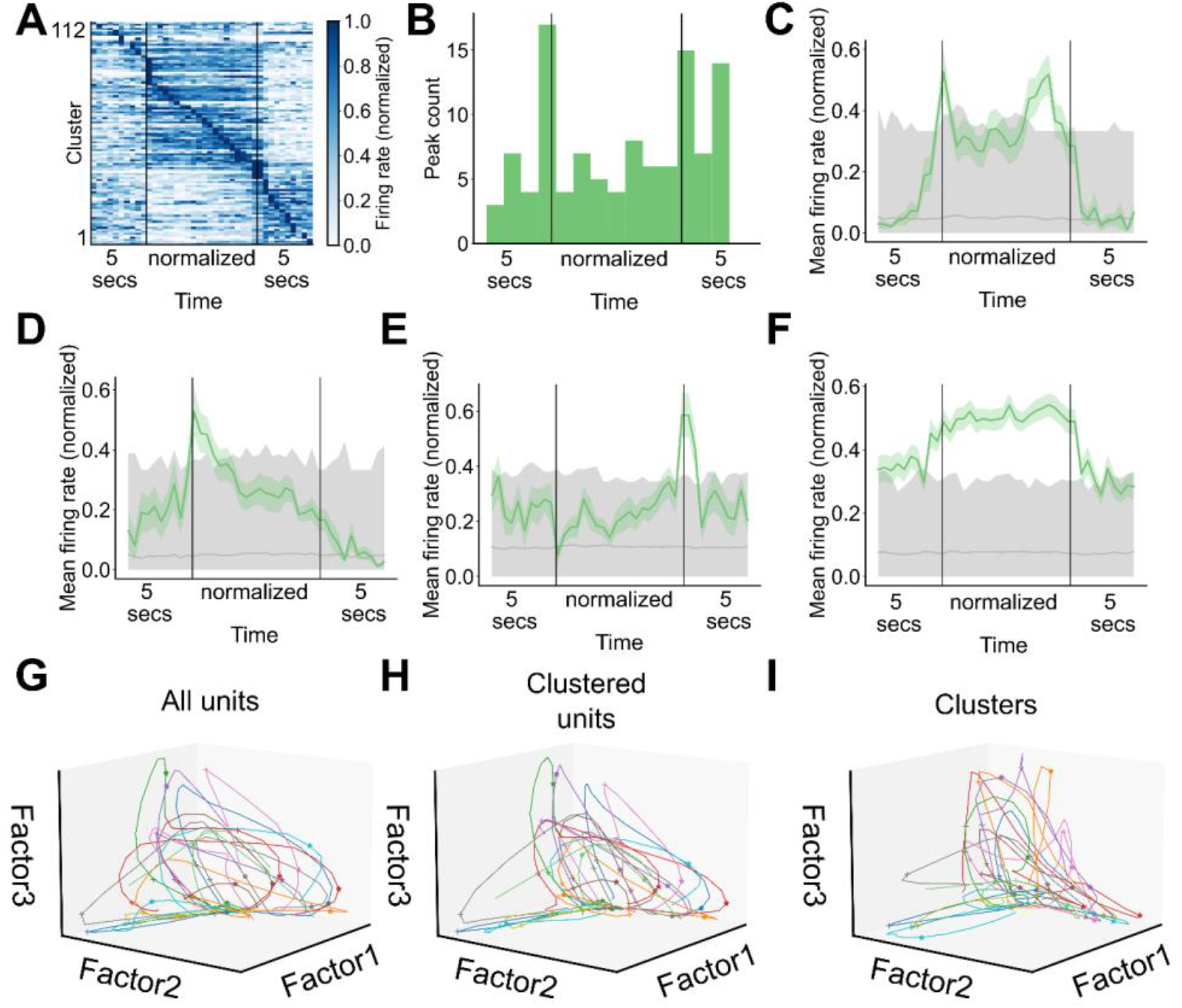
Striatal ensembles encode features of self-grooming. **A**. Heatmap depicting the grooming-aligned average activity of all striatal ensembles. Ensemble averages are normalized to the range from zero to one. Prior to averaging across grooming bouts, ensemble activity during each grooming bout, excluding the 5 seconds before and after grooming, was linearly time warped to a fixed duration. Ensembles are sorted by peak time (112 ensembles, 4 mice, 33 sessions). **B**. Distribution of time at which ensemble average activity peaked (data as in **A**). **C**. Representative example of an ensemble with increased average activity around grooming transitions. Grey region denotes the range of activity for shuffled ensemble activity. Bottom of the range depicts the 2.5^th^ percentile of the shuffled activity, top of the range depicts the 97.5^th^ percentile of the shuffled activity, and grey line depicts the average shuffled activity. **D**. Representative example of an ensemble with increased average activity at the start of grooming (grey region as in **C**). **E**. Representative example of an ensemble with increased average activity at the end of grooming (grey region as in **C**). **F**. Representative example of an ensemble with increased average activity throughout the duration of grooming (grey region as in **C**). **G**. Neural trajectories traced out in factor space by the population of units recorded during all grooming bouts in an example session (65 units, 22 grooming bouts). Colors depict different grooming bouts, pluses denote the start of grooming, and asterisks denote the end of grooming. **H**. Neural trajectories traced out in factor space by the population of clustered units during all grooming bouts in an example session (49 units, 22 grooming bouts). Visualization elements as in **G. I**. Ensemble trajectories traced out in factor space by the striatal ensembles during all grooming bouts in an example session (13 ensembles, 22 grooming bouts). Visualization elements as in **G**.

To contrast the encoding of grooming-related activity in ensembles vs unclustered units, we used nonnegative matrix factorization to perform dimensionality reduction on the neural activity during grooming within a single session. The trajectories traced out by the population activity of all units during all grooming bouts in an example session are shown in **Figure 5G** (65 units). During most grooming bouts the population activity moves out along factor 1, then up along factor 2, and finally returns toward the origin along factor 3, corresponding to population encoding of grooming onset, maintenance, and termination. The neural trajectories computed from unclustered units do not retain the dynamics present in the trajectories from all units or the clustered ones (**Figure 5—figure supplement 1D**, 16 unclustered units; **Figure 5H**, 49 clustered units). Notably, the ensemble trajectories computed from the activity of ensembles retain most of the dynamics seen in the trajectories computed from the activity of single units (**Figure 5I**, 13 ensembles).

**Figure 5–figure supplement 1.**
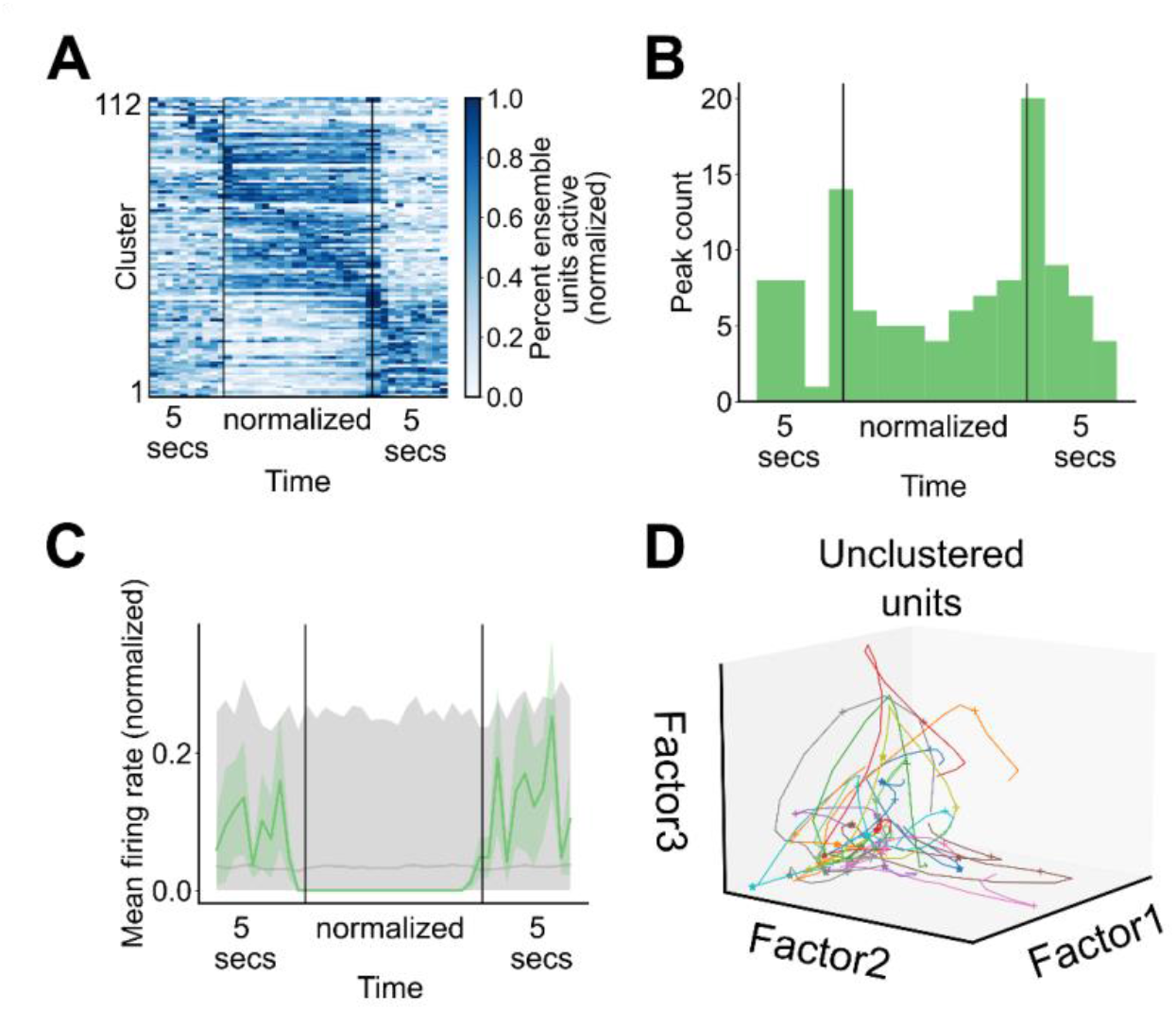
Patterns of cluster engagement. **A**. Heatmap depicting the percent of units within each ensemble that are active during grooming averaged across all grooming bouts in a given session. Prior to averaging across grooming bouts, ensemble activity during each grooming bout was interpolated to a fixed duration, excluding the 5 seconds before and after grooming. Ensembles are in the same order as in **Figure 5A** and data are normalized to the range from zero to one (112 ensembles, 4 mice, 33 sessions). **B**. Distribution of times when the percent of active units in an ensemble peaked (data as in **A**). **C**. Representative example of an ensemble with decreased average activity throughout the duration of grooming (grey region as in **Figure 5C**). **D**. Neural trajectories traced out in factor space by the population of unclustered units during all grooming bouts in an example session (16 units, 22 grooming bouts). Visualization elements as in **Figure 5G**.

## Discussion

Here, we developed a semi-automated approach to identify grooming events in mouse behavioral videos. We recorded population activity in the dorsolateral striatum of freely moving mice and found SPNs and FSIs with activity that encodes the start and end of spontaneous grooming bouts. Previous studies reported changes in striatal single unit activity around the start and end of naturalistic^42,43,52^ and learned behaviors^24,53,54^. Our single unit data elaborate the changes in striatal unit activity around the transitions of a naturalistic behavior. Single unit responses were heterogenous and included units with changes in activity at the start or end of grooming, as well as units that were active or silenced throughout the duration of grooming. We identified striatal ensembles with units that were more correlated during grooming than during the entire session. These striatal ensembles encoded grooming start time, end time, and bout duration. Single session trajectories computed from ensemble activity retain most of the dynamics present in trajectories computed from all units in the session.

The distribution of pairwise distances between units within an ensemble is shifted towards smaller distances, compared to the distribution of distances for units that do not belong to the same ensemble. This is consistent with striatal ensembles detected using imaging techniques^30,38,39^ although our identified ensembles are less spatially compact than those previously reported. Notably, however, striatal 2-photon imaging approaches and microendoscopic imaging of genetically encoded calcium indicators yield a recording area that extends along the medial-lateral and anterior posterior axes (*i*.*e*., horizontally), whereas our four-shank electrophysiological recordings yield a recording field that extends along the anterior-posterior and dorsal-ventral axes (*i*.*e*., vertically), potentially accounting for this difference in findings. Together, the ensemble distances recorded via both methods suggest the possibility that striatal ensembles are isotropically organized, which is well-aligned with studies of cortical and thalamic projection patterns to the striatum^55–57^.

While the ensembles we characterized were identified during grooming behavior, it is possible that they are not grooming-specific. One possibility for the functional organization of striatal ensembles is that ensemble membership is stable across different behaviors. Then, ensemble activation encodes some common movement motif or state that is present across multiple behavioral settings. However, the units within our identified striatal ensembles were more correlated during grooming than during the entire recording session. Our data therefore support an alternative to the stable membership hypothesis, wherein individual units can be members of multiple ensembles, such that when a given unit is active, the behavior the animal is performing can only be determined by looking at the population level. For example, a set of units might be members of a single ensemble that encodes the start of grooming, but each unit could also be part of different ensembles that encode aspects of other natural behaviors, such as eating or walking. This ‘mixed selectivity’ is broadly consistent with previously reported data on striatal activity patterns and cluster memberships that are not conserved across divergent behavioral motifs^30,39^.

Striatal FSIs are parvalbumin-containing GABAergic interneurons that make up approximately 1% of striatal neurons^33,58,59^. FSIs receive direct cortical input, form inhibitory synapses onto SPNs^33,60,61^, and are interconnected amongst themselves via gap junctions on their dendrites^62^. Previous studies found uncoordinated, idiosyncratic task-related changes in FSI activity^63^, as well as changes in FSI activity that correlated with movement features like velocity^64^, with one study showing grooming-related activity in a small number of recorded putative FSIs^43.^ We recorded units that match spike waveform features and firing rates of FSIs^45,59^ and found them to have grooming related activity. These units displayed decreased or increased activity during grooming, as well as transient activity changes at the transitions into and out of grooming. Consistent with strong connectivity present between FSIs and SPNs^65^, a majority of FSIs (70%) were part of identified ensembles and almost all FSIs were part of ensembles together with SPNs.

Striatal control of movement is mediated via two parallel pathways: the direct and indirect pathway, composed of dSPNs and iSPNs^66–68^. Previous recordings of direct and indirect pathway striatal ensemble activity have found that d- and iSPNs exhibit a similar time course of behavior-related changes in their activity^23,30,38^ and have similar ensemble organization^30^, suggesting mixed pathway membership within ensembles. Indeed, simultaneous recording of direct and indirect pathway SPNs showed that striatal ensembles are composed of d- and iSPNs in equal proportions^39^. These prior data suggest that striatal ensembles encoding self-grooming might be composed of both d- and iSPNs, in similar proportions and with similar encoding of grooming, although this remains to be confirmed.

Within bouts of rodent self-grooming, animals perform highly stereotyped grooming sequences called syntactic grooming^69,70^. This sequence is comprised of four distinctive phases and found in a variety of species^8–10^. A large body of work has established a function for the striatum in the sequential ordering of syntactic grooming phases^41,52,71,72^. On a longer time scale, grooming is one of many behaviors an animal can perform at a given time. Since self-grooming, although broadly important, is not often an immediate concern, grooming is considered a low priority behavior^73^. Thus, the control of grooming has been characterized by a disinhibition model, in which grooming takes place in the time left over by other higher priority behaviors when grooming would be inhibited^74^. Previous studies have demonstrated a striatal role in behavioral sequencing on this longer time scale as well^25,30^. Thus, self-grooming represents an ethologically meaningful behavioral paradigm providing a path to study neural control of an innate, conserved behavior at multiple spatiotemporal scales.

## Materials and Methods

### Subjects

Animals were handled according to protocols approved by the Northwestern University Animal Care and Use Committee (protocol number: IS00009022). Adult male and female C57BL/6J mice (p57-105) were used in this study (The Jackson Laboratory, RRID:IMSR_JAX:000664). All mice were group-housed in a humidity-controlled, ambient temperature facility, with standard feeding, 12 hr. light-dark cycle, and enrichment procedures.

### Electrode implantation

Mice were anesthetized with isoflurane (3% for induction, 1.5-2% for maintenance) and placed on a small animal stereotax frame (David Kopf Instruments, Tujunga, CA). A 4-shank 64-channel silicon electrode (part #A4×16-Poly2-5mm-23s-200-177-H64LP_30mm mounted on a dDrive-m, NeuroNexus Technologies, Ann Arbor, MI) was implanted into the dorsolateral striatum (0.5 mm AP, 2 mm ML, and lowered 0.25 mm) and secured with Vetbond (3M, Maplewood, MN) followed by dental cement (Micron Superior 2, Prevest DenPro, Jammu, India, or C&B Metabond, Parkell, Edgewood, NY). The electrode shanks were aligned with the brain’s anterior-posterior axis. A skull screw was connected to the electrode ground wire and fastened to the skull above the ipsilateral cerebellum. The screw was secured with Vetbond followed by dental cement. Mice were monitored following the surgery to ensure a full recovery and were administered post-operative analgesics. Mice recovered for at least 5 days after implantation.

### Behavioral recording

On the first day of experimentation, mice were acclimated to the experimental arena for 30 minutes. All subsequent experimental sessions were 120 minutes long. The experimental arena was an equilateral triangular arena constructed from transparent acrylic (**Figure 1A**. 12-inch sides and height, 1/8-inch thick. Part #8536K131, McMaster-Carr, Elmhurst, IL) fastened together with clear epoxy (Part #31345, Devcon, Solon, OH). While in the arena, mice were given DietGel (ClearH_2_O, Westbrook, ME) for food and hydration. One mouse was in the arena at a time. The arena was cleaned with 70% ethanol after each session. The arena was enclosed by a dark curtain and illuminated by infrared LEDs. Experiments were conducted during the animal’s active phase.

Mouse behavior was captured with three side-view cameras at 125 fps using a custom fork of the campy Python package^75^ (**Figure 1A**. image size 1440 × 608 pixels. cameras: Part #BFS-U3-16S2M-CS Teledyne FLIR, Wilsonville, OR. Lenses: Part #8595755548 Yohii, China). To synchronize recordings from all cameras, each frame grab was triggered by a TTL pulse sent from a microcontroller (Arduino, Turin, Italy). To facilitate alignment of videos and neural data, each frame grab TTL pulse was also sent to the electrophysiology acquisition board and the video and neural recordings were simultaneously initiated by pressing a button connected to the microcontroller.

### Electrophysiological recording

A 64-channel headstage (part #C3325 Intan Technologies, Los Angeles, CA) was connected to the implanted electrode. In the arena, the headstage was connected to a 12-channel commutator supported by a balance arm (Part # FL-12-C-MICRO-BAL Dragonfly Inc., Ridgeley, WV). Neural recordings were acquired at 30 kHz with an Open Ephys acquisition board^76^ and the Open Ephys GUI^76^ (GUI version 0.5.5, Open Ephys, Atlanta, GA). To facilitate alignment of videos and neural data, the video and neural recordings were simultaneously initiated by pressing a button connected to the microcontroller.

Neural recordings were spike sorted offline using KiloSort3^77^ in MATLAB (MathWorks, Natick, MA). We only considered units labeled as ‘good’ by KiloSort. Additionally, we manually inspected the waveforms of all units labeled as ‘good’ and excluded units with waveform shapes that did not resemble an action potential. Recordings were performed from each animal until the number of ‘good units, as identified by KiloSort, stayed below 15 for 2 consecutive days (27.3 ± 4.3 days since implant).

### Unit classification

All recorded units were classified as either putative striatal projection neurons (SPN), fast-spiking interneurons (FSI), or ‘other’ based on their spike waveform and mean firing rate across the entire session following established criteria^45–47^. Units were classified as putative SPNs (88.2 %) if they had peak width > 150 μs, peak-valley interval > 500 μs, and mean firing rate ≤ 10 Hz. Units were classified as FSIs (3.2 %) if they had peak width ≤ 150 μs, peak-valley interval ≤ 500 μs, and mean firing rate ≥ 0.1 Hz. Units that were not identified as either putative SPNs or FSIs were labeled ‘other’ (8.6 %) and excluded from all analyses.

### Histology

After the recordings, coronal brain sections were obtained from all mice to determine electrode placement. Mice were deeply anesthetized with isoflurane and transcardially perfused with 4% paraformaldehyde (PFA) in 0.1 M phosphate buffered saline (PBS). Brains were post-fixed for 1-2 days and washed in PBS. Brains were then sectioned on a vibratome (Leica Biosystems, Wetzlar, Germany) at 60 μm from frontal cortex to posterior striatum and then dried and cover slipped under glycerol:TBS (9:1) with Hoechst 33342 (2.5 μg/ml, Thermo Fisher Scientific, Waltham, MA). Sections were imaged on an Olympus VS120 slide scanning microscope (Olympus Scientific Solutions Americas, Waltham, MA) with DAPI for localization of cell nuclei and FITC for background fluorescence to enable electrode localization.

### Electrode localization

Coronal slices containing electrode tracts were processed using WholeBrain software package^78^ in R^79^. Histological slices were analyzed and registered to the Allen Mouse Brain Common Coordinate Framework (CCFv3)^80^ as described previously^81^. Striatal sections with electrode tracts were analyzed at 60 μm intervals and frontal lobe cortical sections, which rarely contained electrode tracts, were analyzed at 100-200 μm intervals. A total of 6-18 section images were analyzed per mouse brain. Electrode placement for each brain section was denoted at the most ventral location that the electrode tract was observed. Medial-lateral and dorsal-ventral coordinates registered to the CCFv3 were obtained for each section and plotted on a representative coronal section using the WholeBrain package.

### Grooming identification

We identified mouse grooming bouts in behavioral videos using a semi-automated 4-step procedure (**Figure 1C**). As detailed further below, first, we tracked the mouse limb positions in 2D; second, we triangulated to the 2D limb positions to 3D; third, the 3D limb positions were used to isolate likely grooming times; and fourth, using the likely times of grooming we manually identified grooming bout start and stop times.

For 2D pose estimation, we used DeepLabCut (version 2.2.0.5)^82,83^ to detect 15 keypoint positions (snout, both paws, wrists, elbows, shoulders, eyes, ears, and hind paws) on the frames from each camera. We labeled the 15 keypoints in 1,627 frames from 14 videos, 3 animals, and all 3 cameras. We trained a ResNet-50-based neural network^84,85^ with default parameters on 90% of the labeled frames for 300,000 iterations. We validated with 1 shuffle and got a train error of 3.88 pixels and test error of 6.64 pixels (image size: 1440 × 608 pixels). We used a p-cutoff of 0.6 to condition the x and y coordinates for future analyses. With the p-cutoff, the train error was 3.27 pixels and test error 5.06 pixels. This network was then used to analyze all videos from the same experimental setting.

For 3D triangulation of the 2D poses we relied on Anipose^86^. We calibrated our 3 cameras with a 3-minute video of a ChArUco board throughout the camera fields of view (125 FPS. Image size 1440 × 608 pixels). The ChArUco board was printed on paper and taped onto a stiff plastic board. The ChArUco pattern had 7×10 squares containing 4-bit markers and a dictionary size of 50 markers. Before triangulation, we applied a Viterbi filter to the 2D poses with F = 12 frames. The triangulation was performed via an optimization constrained on the distance between the two eyes and the distance between each eye and the snout. We chose scale_smooth = 5, scale_length = 2, and score_threshold = 0.3.

To isolate times when the mouse was likely grooming, we identified frames where the mouse was rearing, and its paws were near its head. To identify frames where the mouse was rearing, we set thresholds on the snout height and the distance between the midpoint of the eyes and the midpoint of the hindlimbs. To identify frames where the mouse’s paws were near its head, we set thresholds on the distance between the midpoint of the paws and the snout, the distance between the midpoint of the paws and the midpoint of the eyes, and height of the midpoint of the paws. Per-frame labels within 1.2 seconds were merged to form predictions over a window. Predicted grooming bouts less than 2 seconds long were discarded. Parameter values were evaluated by generating predictions for a set of videos and viewing the predictions aligned to the video in BENTO^87^.

Grooming bout start and stop times were manually refined by 4 trained annotators, with each annotator labeling a different subset of videos. Annotators used the predictions from the previous thresholding step to navigate the videos and score grooming behaviors. The videos were annotated in VLC media player^88^ using the Time v3.2^89^ and Speed controller^90^ extensions. To assess inter-annotator reliability, all annotators were also given the same 2-hour session to annotate. The average pairwise Jaccard index (or Intersection over Union) computed on the annotations was 0.76 ± 0.04.

### Identification of units with grooming-related activity

To isolate units with grooming related activity, we first down sampled activity to 2 Hz, by summing spike counts within non-overlapping 500 ms bins. We defined units as having grooming-related activity if their average activity at the start or end of grooming was 2 standard deviations above their average activity during a 3 second-long, grooming-free baseline period. For activity aligned to groom start, we used the activity from 5 to 2 seconds before grooming start as the baseline and compared that to the activity from 1 second before until 1 second after the grooming start time. For activity aligned to groom end, the activity from 2 to 5 seconds after grooming was chosen as the baseline and was compared to the average activity from 1 second before until 1 second after groom end. These selection criteria were only used to identify neurons to serve as examples in **Figure 2**; all subsequent analyses were performed on the full dataset.

### Characterization of trial-averaged grooming responses

We characterized the grooming responses exhibited by the recorded units by computing each unit’s event-triggered average response in a ± 5-second window relative to the start and (separately) end of grooming. Responses were down sampled to 4 Hz, by summing spike counts within non-overlapping 250 ms bins. To simplify interpretation, we restricted averaging to grooming bouts that were ‘well-isolated.’ Specifically, because we included the 5 seconds before and after each bout in this analysis, grooming bouts that started within 10 seconds of the end of the previous bout were not included in this analysis. An exception to this is instances with less than 3 seconds between the end of one grooming bout and the start of the next, in which case we merged these two annotations into a single bout.

To better visualize and interpret grooming-associated activity we next performed principal component analysis (PCA)^48^ on the set of event-triggered averages. Specifically, we performed PCA separately on SPN activity aligned to groom start, SPN activity aligned to groom end, FSI activity aligned to groom start, and FSI activity aligned to groom end. For each cell type and condition combination, we concatenated the associated units to form a units-by-time matrix, relative to groom start/stop. The response of each unit in this matrix was then Z-scored, after which PCA was used to extract the first two principal components.

To group units into ‘response types’ for visualization, we examined the weight (or ‘score’) for each unit’s contribution to the largest and second-largest principal component. Units for which the absolute value of this weight was greater for the first principal component than the second formed the “PC1” response group, while those with greater-magnitude weights for the second principal component formed the “PC2” response group. Finally, we sorted the units within each group according to their contribution to the first (PC1) or second (PC2) principal component of the dataset (weight ^*^ activity).

### Grooming-associated striatal ensemble identification

To identify grooming-associated striatal ensembles within a given session, we took the neurons-by-time matrix of recorded spiking from the full two-hour recording session and excluded all frames that did not occur during grooming or within a 5-second window before or after grooming. Due to the low firing rate of SPNs, it was rare to find units that were co-active at high sampling rates. Therefore, to instead identify units that were frequently active at around the same time, we binned spike counts at a sampling rate of 0.667 Hz (bin size of 1.5 seconds). The spike counts for each unit were then normalized to between zero and one, and further smoothed by convolving with a Gaussian filter with a standard deviation of 3 seconds.

Using this smoothed and coarsened estimate of cell activity, we next applied meta-*k*-means^51^ to identify sets of cells that were often co-active. We chose to use meta-*k-*means for clustering, because it does not force all units in a session to be assigned to a cluster, and it allows for the number of ensembles to be greater or less than the initial choice of *k*. Briefly, meta-*k*-means employs repeated runs of standard *k*-means clustering to identify groups of units that are consistently clustered together. As with standard *k*-means, meta-*k*-means requires the user specify an initial number of clusters *k*, which we set to be the square root of the number of units in the given session, as in Barbera et al^38^. We then ran 1,000 repeats of the *k*-means clustering algorithm on the matrix of filtered grooming-related activity described above, with units as variables and time bins as observations. Cluster centroids for each repeat were initialized using the greedy *k*-means++ algorithm^91^. As in prior work^30,38,92^, units that were assigned to the same cluster in >80% of *k*-means runs were considered to be part of the same meta-*k*-means cluster.

When merging clusters as part of the meta-*k*-means algorithm, we used the silhouette score^93^ to evaluate the outcome of a potential cluster merge. To ensure adequate sample size for downstream analyses, ensemble identification was restricted to sessions with at least 30 SPNs and FSIs.

### Statistical analysis of striatal ensembles

To compute a statistical estimate on the number of ensembles present in each session we used an eigenvalue based statistical method from Peyrache at al^49,50^. Briefly, if two neurons have correlated firing, then we expect to observe common fluctuations in their spiking activity during recording. This can be visualized by plotting the firing rates of the two neurons against each other, in the form of a direction in neural activity space along which observations tend to be distributed. Conversely, without any correlated firing, the variance in the dataset will be roughly equally distributed in all directions, and one would not expect to see structure when plotting the activity of two cells against each other. These relationships hold for any n-dimensional set of neurons, where structure emerges from correlated activity. The eigenvectors of the covariance matrix of a dataset represent the directions of maximally shared variance, thereby capturing the correlations present in the data. Each eigenvalue quantifies the variance of the data along the axis defined by its corresponding eigenvector. Thus, in a matrix of neurons-by-time, the eigenvalues of its covariance matrix capture the correlations among neurons (*i*.*e*., ensembles) and comparing the eigenvalues to those computed from the covariance matrix of a shuffled version of the data provides a measure of the number of ensembles present.

To apply this method, we first binned spike counts at the same sampling rate as for clustering (0.667 Hz, bin size of 1.5 seconds) and Z-scored the activity of each neuron. Next, we shuffled the time bins for each neuron independently, and computed the maximal eigenvalue of the covariance matrix of the shuffled data. We repeated this 5,000 times to form a null distribution and counted the number of eigenvalues from the covariance matrix of the actual data that were above the 99^th^ percentile of this null distribution.

### Time-normalized grooming-related activity

To compare ensemble average activity during grooming across grooming bouts of varying duration, we linearly temporally rescaled the activity of each unit during each grooming bout to a single fixed length. We chose the length for the interpolated values from the distribution of grooming bout durations from all clustered sessions. To avoid ‘blurring’ activity associated with the start and end of grooming bouts, the activity during the 5 seconds before and after each bout was not interpolated.

### Bootstrap significance testing of grooming-related ensemble activity

To test for periods during grooming when the grooming ensemble average activity was significantly higher or lower than chance, we computed the mean, 2.5^th^, and 97.5^th^ percentiles of the distribution of randomly sampled ensemble activity. To generate this distribution, we computed the population-average ensemble activity during 1,000 random duration windows of time sampled throughout the 2-hour recording session. Start times were sampled uniformly from 5 seconds into the recording session until 5 seconds before the end of the session. Window durations were sampled from the distribution of observed grooming bout durations across all sessions.

### Neural trajectories

To visualize neural trajectories, neural activity during grooming was decomposed onto 3 components using non-negative matrix factorization^94^. For the neural trajectories, we decomposed a matrix containing each unit’s activity during all grooming bouts within that session. For each unit, we concatenated its activity during all grooming bouts including the five seconds before and after each bout. For the ensemble trajectories, we averaged the activity of all units within each ensemble and concatenated each ensemble’s activity during all grooming bouts including 5 seconds before and after each bout. We initialized using Nonnegative Double Singular Value Decomposition with zeros filled with the average neural activity and minimized the Frobenius norm of the loss.

### Statistical methods

All statistical tests were two-sided. Statistical significance was set to p = 0.05. Summary data in all figures are reported as mean ± SEM.

### Software

All custom software was written in Python^95^ unless stated otherwise. In addition to those mentioned elsewhere, we used the following Python packages: numpy^96^, scipy^97^, matplotlib^98^, scikit-learn^99^, and _pandas_^100,101^.

## Acknowledgements

The authors would like to thank Lindsey Butler for mouse colony management.

This research was supported in part through the computational resources and staff contributions provided for the Quest high performance computing facility at Northwestern University which is jointly supported by the Office of the Provost, the Office for Research, and Northwestern University Information Technology.

Data visualization colors were chosen from ColorBrewer^102^.

## Author contributions

Conceptualization: SM, AK, YK. Methodology: SM. Software: SM. Validation: SM, YK. Formal Analysis: SM. Investigation: SM, MAM, FHM, HY, SNF, EC. Resources: SM, AK, YK. Data Curation: SM. Writing - Original Draft Preparation: SM, YK. Writing - Review and Editing: SM, MAM, FHM, HY, SNF, EC, AK, YK. Visualization: SM, SNF. Supervision: AK, YK. Project Administration: SM, AK, YK. Funding Acquisition: AK, YK.

The following authors contributed equally, and their names were ordered alphabetically in the author list: MAM, FHM, and HY.

## Code availability

Upon publication, code will be available in a public repository on our lab GitHub page (https://github.com/kozorovitskiylaboratory).

